# *Leptin* deficient rats develop nonalcoholic steatohepatitis with unique disease progression

**DOI:** 10.1101/594978

**Authors:** Ping Lu, Guang Yang, Wen He, Wanwan Wu, Lingbin Qi, Shijun Shen, Junhua Rao, Guoping Fan, Zhigang Xue, Peng Zhang, Cizhong Jiang, Xianmin Zhu

**Affiliations:** Tongji University, School of Medicine, Shanghai Pulmonary Hospital, Shanghai 200433, China; Translational Center for Stem Cell Research, Tongji Hospital, Department of Regenerative Medicine, Tongji University School of Medicine, Shanghai 200065, China; Institute of Translational Research, Tongji Hospital, the School of Life Sciences and Technology, Shanghai Key Laboratory of Signaling and Disease Research, Tongji University, Shanghai 200092, China; Tongji University, School of Life Sciences and Technology, Shanghai 200092, China; Guangdong Key Laboratory of Animal Conservation and Resource Utilization, Guangdong Public Laboratory of Wild Animal Conservation and Utilization, Guangdong Institute of Applied Biological Resources, 105 Xingang Rd. West, Guangzhou, Guangdong 510260, China; Department of Human Genetics, David Geffen School of Medicine, University of California Los Angeles, Los Angeles CA 90095, USA; The Research Center of Stem Cells and Ageing, Tsingtao Advanced Research Institute, Tongji University, 5 Nanhai Branch Road, Tsingdao 266071, China

**Keywords:** nonalcoholic fatty liver disease, transcriptome, triglyceride biosynthesis, hepatic inflammation, natural history

## Abstract

Nonalcoholic steatohepatitis (NASH) is an aggressive liver disease threatening public health, however its natural history is poorly understood. Unlike *ob/ob* mice, *Lep^∆I14/∆I14^* rats develop unique NASH phenotype with steatosis, lymphocyte infiltration and ballooning after postnatal week 16. Using *Lep^∆I14/∆I14^* rats as NASH model, we studied the natural history of NASH progression by performing an integrated analysis of hepatic transcriptome from postnatal week 4 to 48. *Leptin* deficiency results in a robust increase in expression of genes encoding 9 rate-limiting enzymes in lipid metabolism such as ACC and FASN. However, genes in positive regulation of inflammatory response are highly expressed at week 16 and then remain the steady elevated expression till week 48. The high expression of cytokines and chemokines including CCL2, TNFα, IL6 and IL1β is correlated with the phosphorylation of several key molecules in pathways such as JNK and NF-κB. Meanwhile, we observed cell infiltration of MPO^+^ neutrophils, CD8^+^ T cells, CD68^+^ hepatic macrophages and CCR2^+^ inflammatory monocyte-derived macrophages, together with macrophage polarization from M2 to M1. Importantly, *Lep^∆I14/∆I14^* rats share more homologous genes with NASH patients than previously established mouse models and crab eating monkeys with spontaneous hepatic steatosis. Transcriptomic analysis showed that many drug targets in clinical trials can be evaluated in *Lep^∆I14/∆I14^* rats.

**Conclusion:** We characterize *Lep^∆I14/∆I14^* rats as a unique NASH model by performing a long-term (i.e., 4 to 48 postnatal weeks) integrated transcriptomic analysis. This work reveal the temporal dynamics of hepatic gene expression in lipid metabolism and inflammation, and shed light on understanding the natural history of NASH in human beings.

## Introduction

Nonalcoholic fatty liver disease (NAFLD) is the most common liver disease worldwide, affecting approximately 24% of the population.(1) Depending on the presence of inflammation, it is further categorized into nonalcoholic fatty liver (NAFL) and nonalcoholic steatohepatitis (NASH).(2) NAFLD, especially NASH, will eventually develop into liver fibrosis, cirrhosis, and hepatocellular carcinoma (HCC) in some cases.(3) Historically, the severity of NAFLD can be assessed by many diagnostic modalities, including examination of demographic record, serum parameters, histopathological characteristics of liver biopsies and the emerging non-invasive methods such as magnetic resonance imaging-proton density fat fraction (MRI-PDFF) and magnetic resonance elastography (MRE).(4) However, the severity level only provides snapshot of a certain stage during the temporal progression of the disease. The natural history of NAFLD in human beings is still poorly understood due to its slow progression (5) and ethical issues.

LEPTIN is an adipokine secreted by white adipocyte tissues. Binding to its receptor (LEPR), LEPTIN can precisely regulate food intake, glucose and lipid metabolism, energy homeostasis, immune system, reproduction, etc.(6, 7) *Leptin* deficient *ob/ob* mice (8) have been discovered decades before the gene *Leptin* was characterized by Friedman’s laboratory (6). *ob/ob* mice are widely used as NAFLD model to mirror the human disease with obesity, hyperglycaemia, and hyperinsulinaemia.(9) But it should be noted that *ob/ob* mice do not develop spontaneous NASH.(10, 11) Therefore, little is known about LEPTIN function during NASH progression although it is a pro-inflammatory factor (12) and required for the development of fibrosis in NASH (13). We previously generated *Leptin* deficient (*Lep^∆I14/∆I14^*) rats by CRISPR technology.(14) Interestingly, we found that *Lep^∆I14/∆I14^* rats display distinctive NASH phenotypes, which needs to be further characterized.

To study the pathogenesis and progression of NAFLD, many animal models were generated by genetic manipulation and/or dietary induction.(9–11, 15) However, the previous studies mainly focused on the endpoint when certain histopathological phenotype was observed, and neglected the temporal progression of NAFLD. Therefore, we used *Lep^∆I14/∆I14^* rats as a NASH model and performed a long-term (i.e., 4 to 48 postnatal weeks) integrated transcriptomic analysis, aiming to reveal the temporal dynamics of hepatic gene expression due to LEPTIN deficiency and shed light on understanding the natural history of NASH in human beings.

## Materials and Methods

### Animals

Rats and mice were kept in a 12-h light, 12-h dark cycle with *ad libitum* access to regular chow food and water. The controls were littermates. B6/JNju-Lep^em1Cd25^/Nju mice were purchased from Nanjing Biomedical Research Institute of Nanjing University (Cat#T001461). All the protocols were in accordance with the guidelines of Tongji University’s Committee on Animal Care and Use. All the experimental procedures as described below were approved by the animal experiment administration committee of Tongji University (# TJLAC-016-021). PharmaLegacy Laboratories (Shanghai) Co., Ltd performed a pre-screening of *Macaca fascicularis* for spontaneous NAFLD. All procedures related to *Macaca fascicularis* were approved by PharmaLegacy Laboratories’ IACUC and followed the guidelines of Association for Assessment and Accreditation of Laboratory Animal Care (AAALAC).

### Serum biochemical analyses

The animals were fasted for 16 h and blood was collected and left to clot for 1 h at room temperature. Serum was obtained after centrifugation. Serum alanine aminotransferase (ALT) and aspartate aminotransferase (AST) were determined by using the spectrometric kits (Sigma Diagnostics). Triglyceride, high density lipoprotein cholesterol (HDL-C) and low density lipoprotein cholesterol (LDL-C) were examined by Adicon Central Lab. Serum IL1β was detected by Elisa kit (Boster Biological Technology).

### Histological analyses

Liver samples of the animals were collected and fixed according to the routine methods for hematoxylin and eosin (HE) staining, Oil red O staining, Sirius red staining, and IHC respectively. The detailed information of the antibodies for IHC was summarized in table S1.

### RT-PCR

Total RNA was extracted from snap-frozen liver samples using TRIzol (Life Technologies) according to the manufacturer’s instructions. The complementary DNA was generated from 2 µg total RNA by RevertAid First Strand cDNA Synthesis Kit (Thermo Scientific). Quantitative PCR (qPCR) was performed using TB GreenTM Premix Ex TaqTM II (TaKaRa, RR820A) on QuantStudio 7 Flex Real-Time PCR System (Applied Biosystems). PCR primers were designed to specifically amplify genes of interest (table S2). Quantitative polymerase chain reaction (qPCR) was performed with β-Tubulin as internal control.

### Western blot

Total protein was extracted from snap-frozen liver samples using RIPA lysis buffer (EpiZyme, PC104) with protease inhibitors (EpiZyme, GRF101) and phosphatase inhibitors (EpiZyme, GRF102). Western blot was performed following the routine method. The antibodies were listed in table S1. All western blots were normalized to β-Tubulin for each sample.

### RNA-seq and bioinformatics

Total RNA was extracted from snap-frozen liver samples using TRIzol (Life Technologies) according to the manufacturer’s instructions. RNA purity was checked using the Nano Photometer spectrophotometer (IMPLEN). RNA concentration was measured using Qubit RNA Assay Kit in Qubit 2.0 Flurometer (Life Technologies). RNA integrity was assessed using the RNA Nano 6000 Assay Kit of the Bioanalyzer 2100 system (Agilent Technologies). Library construction was done using NEBNext Ultra Directional RNA Library Prep Kit for Illumina (NEB) following manufacturer’s protocols. After cluster generation (TruSeq PE Cluster Kit v3-cBot-HS [Illumia]), the libraries were sequenced on an Illumina Hiseq X platform and 150 bp paired-end reads were generated in the facilities of Novogene.

RNA-seq data analysis. Using Hisat2 (v2.1.0), the rat RNA-seq reads were mapped to the rn6 reference genome (RefSeq) and the crab-eating macaque RNA-seq reads were mapped to *Macaca fascicularis* (v5.0.91) reference genome (Ensembl). For transcriptome quantification, we used featureCounts (v1.6.1) to generate read counts for all genes. Raw counts were normalized by DESeq2 (v1.20.0) using rlog method and normalized counts of replicates were averaged. The differentially expressed genes between two samples were identified by DESeq2 (v1.20.0) with a Benjamini-Hochberg-adjusted p value < 0.05 and a fold change > 1.5.

Gene ontology (GO) analysis. We performed functional annotation of gene sets using the Database for Annotation, Visualization and Integrated Discovery (DAVID) Bioinformatics Resource. P values for clusters were plotted to show the significance of functional clusters.

Time series analysis of fat accumulation and inflammation geneset. Genes represent each feature were collected as described in the main text. Their gene expression levels were normalized by Z-score and presented as scatter plots.

Species comparison analysis. Homologous genes of four species (Human, crab-eating macaque, rat, mouse) were downloaded from the Ensembl genome browser Release 91 (http://dec2017.archive.ensembl.org/index.html). The significantly differentially expressed genes and pathways in human beings with healthy obesity (HO), NAFLD, and NASH were retained from the previous study *(17)*.

Accession numbers. The accession number for the rat RNA-seq data generated in this study is GSE124002.

### Statistical Analyses

At each time point, the data from *Lep^∆I14/∆I14^* rats were compared to the WT controls by t test. P value of <0.05 was defined as the level of significance, and all graphs and analyses were performed using GraphPad Prism (version 6; GraphPad Software Inc., La Jolla, CA).

## Results

### *Leptin* deficiency leads to NASH phenotypes in rats

*Lep^∆I14/∆I14^* rats share many similar phenotypes with *ob/ob* mice, such as obesity, hyperglycaemia, hyperinsulinemia, NAFLD, and infertility.(14) Our previous histopathological examination showed that NAFLD starts at postnatal week 8.(14) Consistently, body weight and relative liver weight of *Lep^∆I14/∆I14^* rats were significantly higher than those of their wild-type (WT) littermates after 8 postnatal weeks (fig. S1A). We also observed the increase in serum parameters such as triglyceride, high density lipoprotein cholesterol (HDL-C) and low density lipoprotein cholesterol (LDL-C) (fig. S1A). Interestingly, alanine aminotransferase (ALT) and aspartate aminotransferase (AST) levels did not follow a trend of rectilinear increase but have an inflection point at postnatal week 16 (fig. S1A), suggesting that NASH related inflammation develops but is restricted at certain level afterwards. To further characterize NASH phenotypes in *Lep^∆I14/∆I14^* rats, we did hematoxylin-eosin (HE) and Oil Red O staining of liver sections prepared at different time points during disease progression (Fig. 1A). Simple steatosis was developed after week 8, however, it was at week 16 that emerged the stereotypic NASH phenotypes including steatosis, lymphocyte infiltration and ballooning (Fig. 1B-D). MPO^+^ neutrophil infiltration started to be observed at week 16 (Fig. 1E). Using the NASH scoring system (table S3) (16), we found that *Lep^∆I14/∆I14^* rats indeed developed NASH (NASH score≥5) which reached a plateau after week 16 (Fig. 1F). We did not see fibrosis in *Lep^∆I14/∆I14^* rats at any time points through week 48 (fig. S1B).

**Fig. 1.**
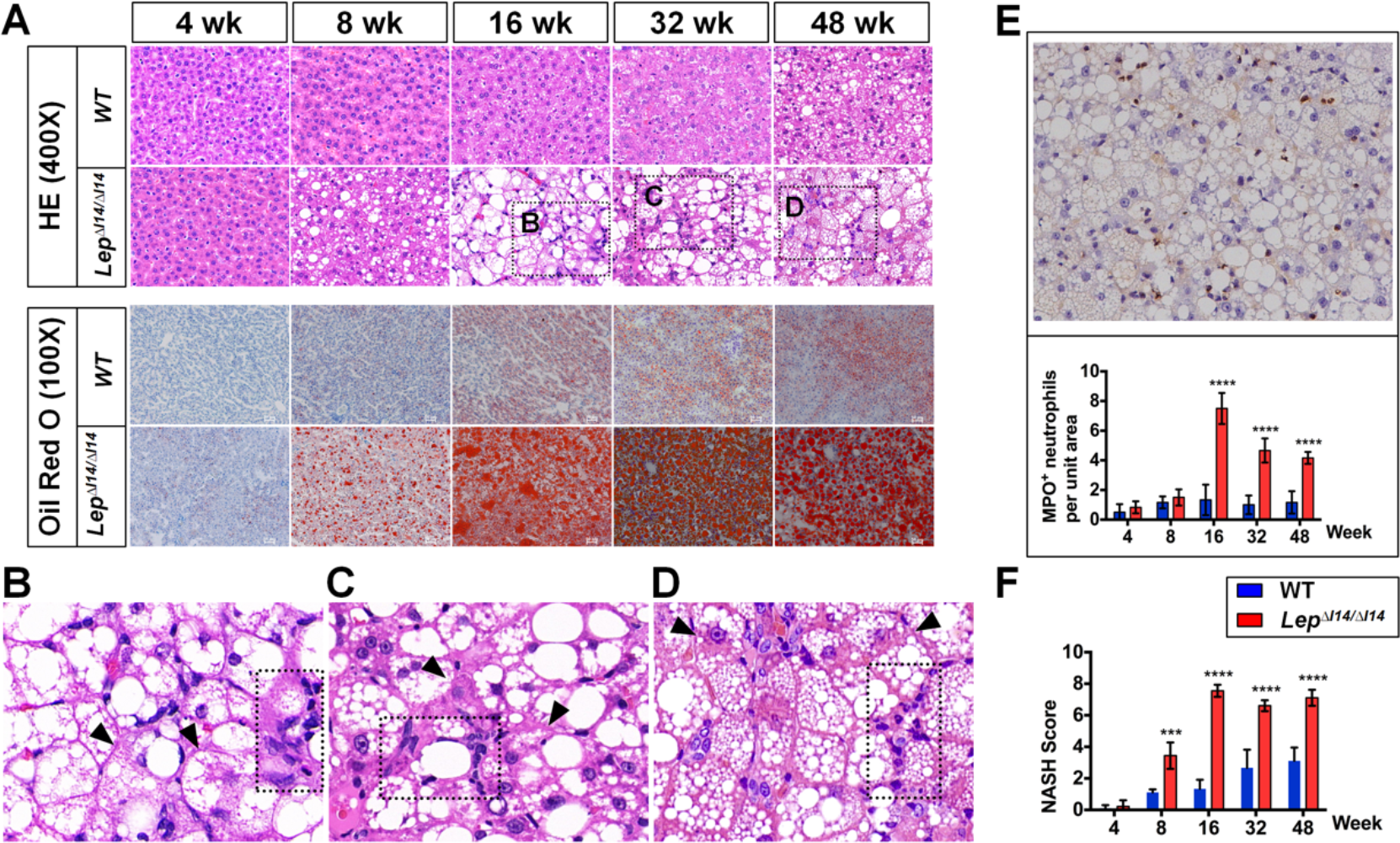
Unique NASH phenotypes of *Lep^∆I14/∆I14^* rats. (A) HE and Oil red O staining of liver sections shows that NASH phenotypes start to be observed at week 16. (B-D) Magnified the field of HE staining in (A) showing NASH phenotype at week 16, 32 and 48 respectively. Arrow heads indicate hepatocyte ballooning; dashed rectangle indicates lymphocyte infiltration. (E) MPO staining shows neutrophil infiltration. Left panel shows a representative picture at week 16, and right panel is the statistical analysis of each time point (n=6). (F) The NASH scores at each time point. ***p<0.001, ****p<0.0001.

As previous research showed that *ob/ob* mice do not develop spontaneous NASH (10, 11), we performed the independent experiments to validate this in our laboratory. Similar to *Lep^∆I14/∆I14^* rats, *Leptin* deficient mice had significantly higher body weight (Fig. 2A), relative liver weight (Fig. 2B), AST (Fig. 2C) and ALT (Fig. 2D) at postnatal week 16. As expected, they had simple steatosis without NASH-like phenotype (Fig. 2E). Compared to the controls, they did not show any difference in immune cell infiltrates including LY6G^+^ neutrophils, F4/80^+^ macrophages, CD11C^+^ M1 macrophages, CD163 M2 macrophages and CCR2^+^ inflammatory monocyte-derived macrophages (Fig. 2F). We did not observe NASH-like phenotype in *Leptin* deficient mice at week 32 (fig. S2). Taken together, *Leptin* deficiency leads to NASH phenotype in rats but not mice.

**Fig. 2.**
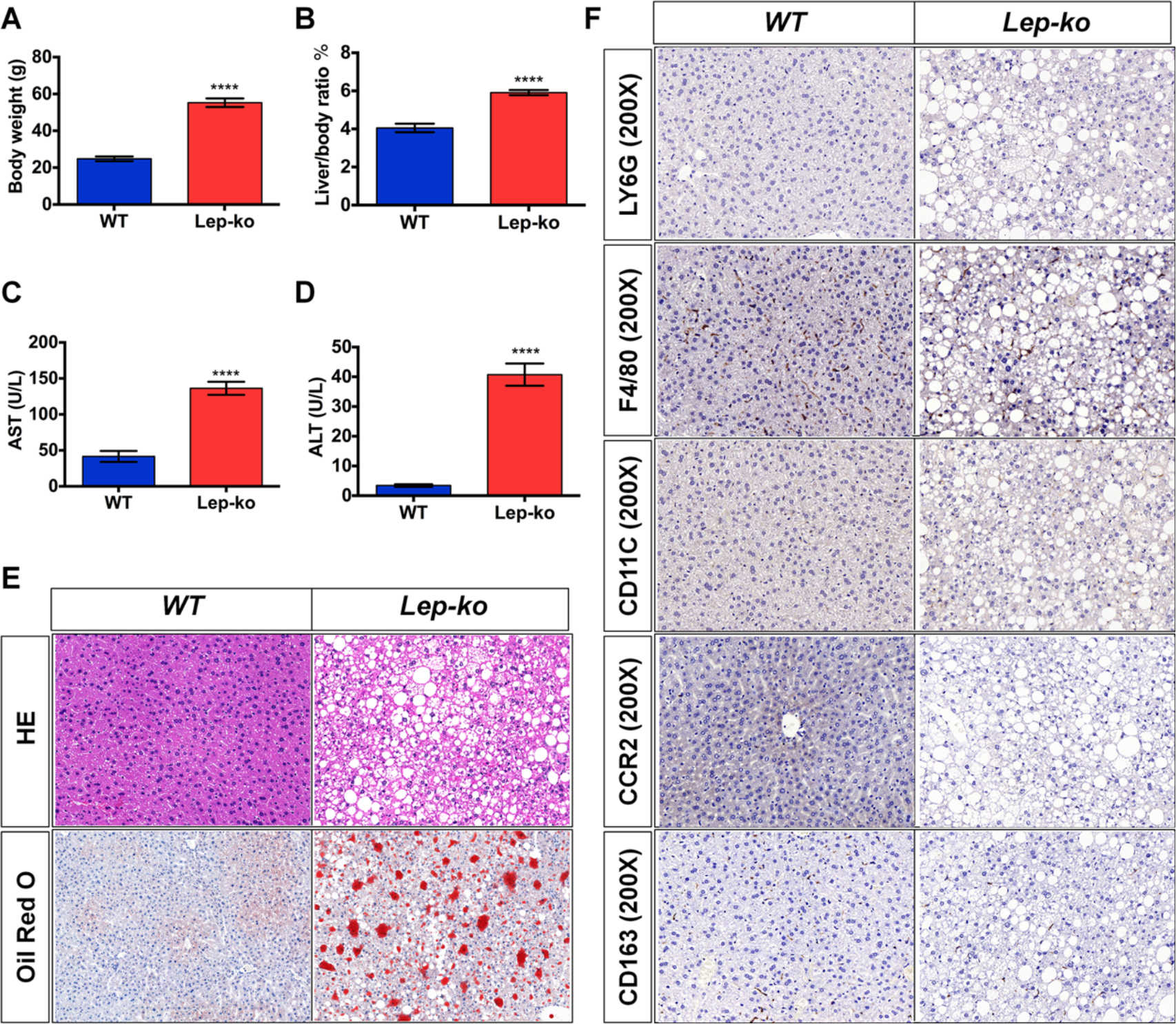
*ob/ob* mice do not develop NASH phenotypes. (A-D) Body weight, liver/body ratio, ALT and AST in *ob/ob* mice at week 16 (n=6). Data are presented as Mean±SD. ****p<0.0001. (E) HE and Oil red O staining showed liver steatosis at week 16. (F) Immune cell infiltration in liver showed no difference between *ob/ob* mice and the WT controls at week 16.

### Integrated transcriptomic analysis show that NASH progression involves distinct gene regulation of lipid metabolism and hepatic inflammation

To understand the disease progression at systemic level, some pioneers have investigated the transcriptome of liver biopsies from NAFLD patients at mild, medium and severe stages.(17, 18) However, the severity levels in different individuals do not reflect the temporal progression of the disease in a particular person. Therefore, we decided to build a more complete picture of NASH progression by using this unique model of *Lep^∆I14/∆I14^* rats. We performed RNA-sequencing (RNA-seq) of the liver samples from WT and *Lep^∆I14/∆I14^* rats at postnatal 4-week, 8-week, 16-week, 32-week and 48-week respectively. Over all, the hepatic transcriptome showed dynamic changes at different time points (Fig. 3A). At week 4, the expression profiles of WT controls and *Lep^∆I14/∆I14^* rats were very similar and cluster together, which is in line with the histopathological data (Fig. 1A) that *Lep^∆I14/∆I14^* rats have not yet developed NAFLD by week 4. From week 8 to week 32, the expression profiles of WT controls and *Lep^∆I14/∆I14^* rats clustered separately, implying a temporal NAFLD progression. It should be noted that the expression profiles of WT controls at week 48 are clustered closely to that of *Lep^∆I14/∆I14^* rats (Fig. 3A), probably due to the aging-related NAFLD in WT controls (Fig. 1A). We then identified the differentially expressed genes (DEGs) between *Lep^∆I14/∆I14^* and WT rats for each time point and annotated their functions with enriched gene ontology (GO) terms (Fig. 3B). The results showed that the up-regulated DEGs at week 4 and 8 were enriched mainly for lipid-metabolism related GO terms (Fig. 3B). However, up-regulated DEGs from week 16 to 48 were enriched for inflammation related GO terms (Fig. 3B), which was in parallel with the NASH phenotypes as previously described in Fig 1. Interestingly, only three up-regulated (Fig. 3C) and eight down-regulated (fig. S3) DEGs were common across all time points. Instead, considerable DEGs were common between adjacent time-points (table S4), implying that NAFLD progresses stepwisely with activation/inactivation of different sets of genes. We then asked if the trend of gene expression related to lipid accumulation and inflammation can represent NASH progression with week 16 as the inflection point between simple steatosis and NASH. To answer this question, we first profiled the dynamic expression of the genes with functions related to lipid metabolism and inflammation respectively (fatty acid biosynthetic process [GO:0006633] and positive regulation of inflammatory response [GO:0050729]). As expected, the inflammation-related genes in *Lep^∆I14/∆I14^* rats started to be expressed significantly higher than WT at week 16 (p<0.05) and then remained the steady elevated expression (p<0.05) till week 48 (Fig 3D). To our surprise, the expression levels of lipid-metabolism related genes have no significant difference between *Lep^∆I14/∆I14^* and WT rats for each time point, and have no precipitous fluctuation across all time points (Fig. 3D). As liver steatosis is morbid accumulation of triglyceride (TAG) in hepatocytes, we selected 9 genes which encode well-known essential enzymes including ACC and FASN for TAG synthesis. The expression of these 9 genes displayed an elevating trend in both WT and *Lep^∆I14/∆I14^* rats, however, was significantly higher in *Lep^∆I14/∆I14^* rats (Fig. 3E). All the 9 genes had a dynamic expression pattern during NASH progression (Fig. 3F), some of which were verified by qRT-PCR (fig. S4A). This suggests that liver steatosis is correlated with the elevated expression of a subgroup of rate-limiting enzymes, which may provide combinatory therapeutic targets for reducing fat accumulation in liver. In conclusion, our data support that NASH progression is a continuous process which coordinates distinct gene expression involving in lipid metabolism and inflammation.

**Fig. 3.**
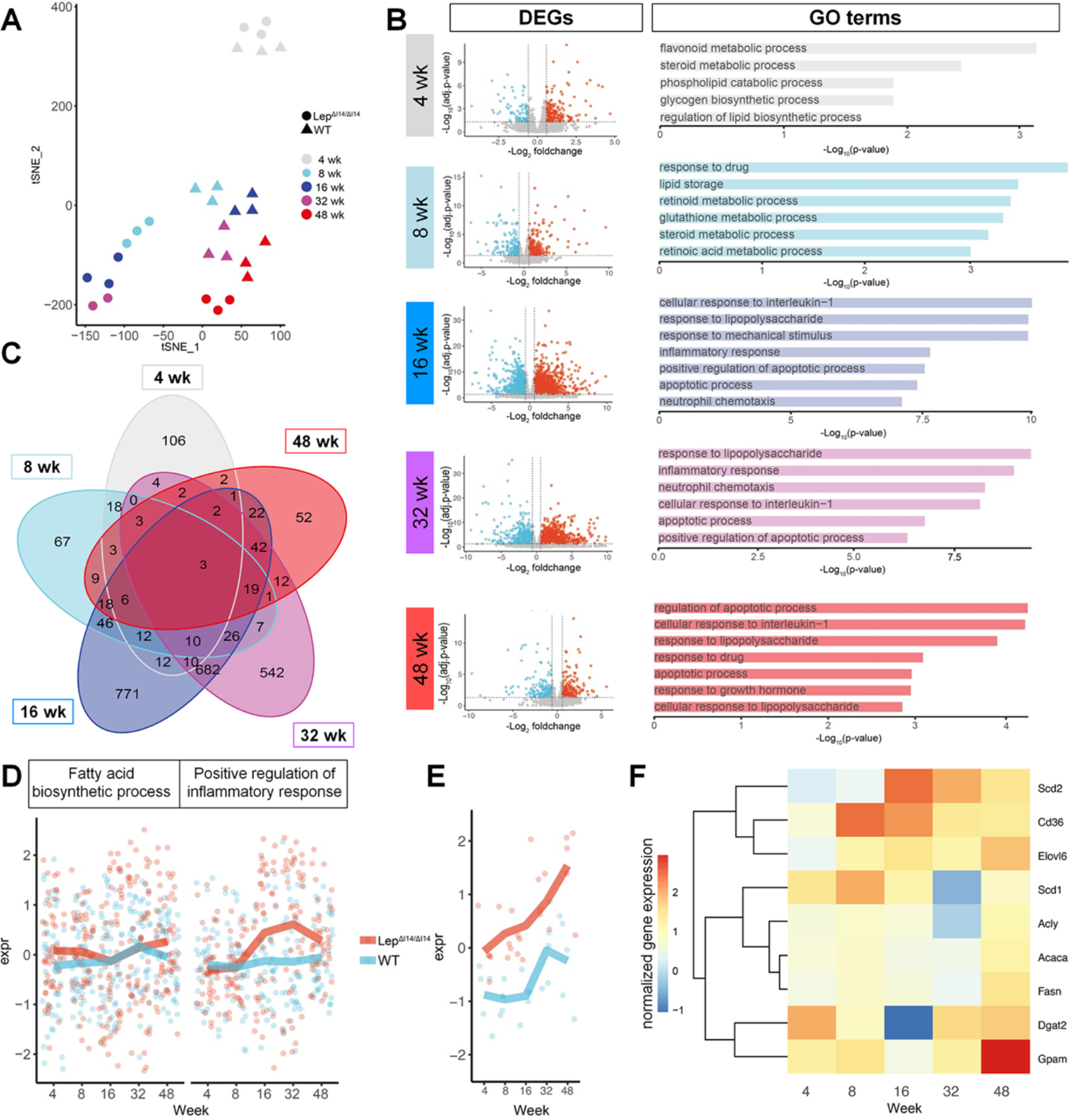
Hepatic transcriptome alteration during NASH progression in *Lep^∆I14/∆I14^* rats. (A) tSNE plot of hepatic gene expression in *Lep^∆I14/∆I14^* rats and the WT controls from postnatal week 4 to 48 (n=3 for samples at each time point except n=2 for *Lep^∆I14/∆I14^* rats at week 32). (B) Volcano plots show the DEGs across different time points. Blue and red dots are down- and up-regulated DEGs, respectively. Bar plots are the GO terms for which the up-regulated DEGs enriched. (C) Venn plot shows the overlap of up-regulated DEGs among different time points. (D) The expression trend of DEGs in GO terms “fatty acid biosynthetic process” (GO:0006633) and “positive regulation of inflammatory response” (GO:0050729). (E) The expression trend of 9 genes encoding rate-limiting enzymes for TAG synthesis. (F) Heatmap showing expression levels of the genes in (E).

### Restricted hepatic inflammation in *Lep^∆I14/∆I14^* rats is determined by the inflammation related genes, pathways and immune cell infiltration

To explain the restricted inflammation after week 16 during NASH progression in *Lep^∆I14/∆I14^* rats, we decided to investigate the inflammation related genes, pathways and immune cell infiltration. As demonstrated above (Fig. 3D), many pro-inflammation genes started to be expressed with a high level at week 16 in *Lep^∆I14/∆I14^* rats (Fig. 4A). We also examined the expression of several inflammation related genes including *Ccl2*, *Tnfα* and *Il6* by qRT-PCR (Fig. 4B) and/or Western blot (fig. S4B and C), and found that their expression levels indeed peaked at week 16. Meanwhile, serum IL1β level was elevated at week 16 and remained at a steady level afterwards (Fig. 4C). It’s well known that inflammatory processes are mainly regulated by conserved signaling pathways through the phosphorylation of several key molecules such as JNK and NF-κB (19, 20). So we examined the expression and post-translational modification of these molecules at different time points by Western blot (Fig. 4D and E). We found that the activated (phosphorylated) form of JNK (Fig. 4D) was markedly higher after week 16 in *Lep^∆I14/∆I14^* rats compared to the WT controls. The elevation of phosphorylated P65 was observed as soon as week 8 (Fig. 4D), which is in line with the fact that NF-κB mediated signaling is required for the activation of pro-inflammation gene expression.

**Fig. 4.**
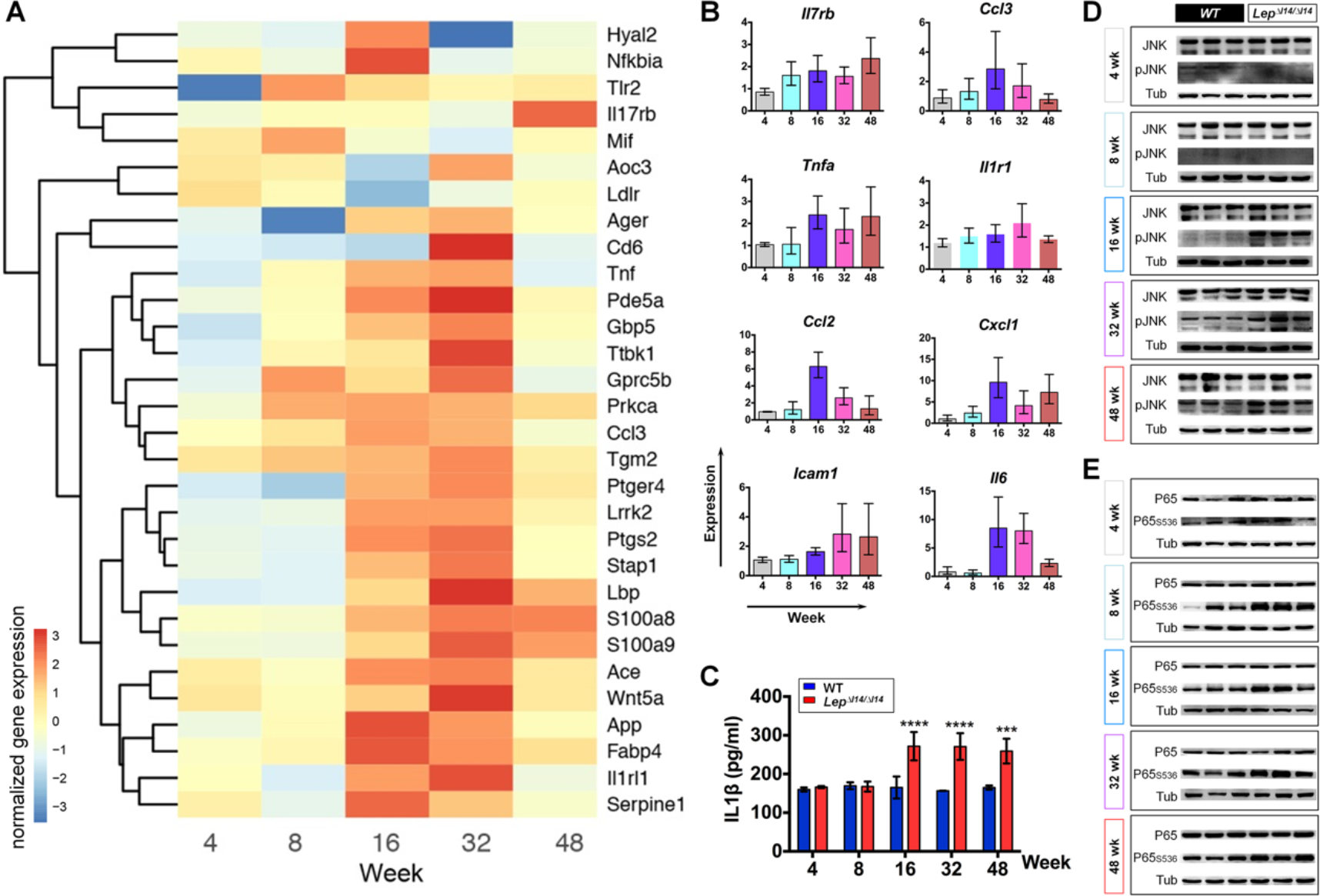
The inflammation related genes and pathways during NASH progression in *Lep^∆I14/∆I14^* rats. (A) Heatmap of the relative gene expression related to inflammation in *Lep^∆I14/∆I14^* rats. (B) qRT-PCR verification of the expression of some inflammation related genes (n=3). Shown is the relative expression using *Actb* as internal control. Data are presented as Mean±SD. (C) Serum IL1β levels at different time points (n=3). Data are presented as Mean±SD. ***p<0.001, ****p<0.0001. (D) and (E) Western blot shows the levels of phosphorylated JNK (D) and p65 (E) at different time points (n=3).

Macrophage polarization of pro-inflammatory M1 (classically activated macrophages) and anti-inflammatory M2 (alternatively activated macrophages) also contributes to liver inflammation and injury (21). Interestingly, we found that M2 related gene *Arg1* was downregulated in the livers of *Lep^∆I14/∆I14^* rats at week 16, while M1 related gene *Inos* (*Nos2*) was upregulated, which was validated by qRT-PCR (Fig. 5A). At protein level, M2 marker CD163 was reduced at week 16 (Fig. 5B), while M1 marker CD11B (22, 23) was increased (Fig. 5C). Since inflammation in liver involves infiltration of distinct types of immune cells (24, 25), this suggested that NASH progression in *Lep^∆I14/∆I14^* rats may also involve immune cell infiltration. Indeed, we observed an increase of CD8^+^ T cells and CD68^+^ hepatic macrophages (Fig. 5D) in addition to MPO^+^ neutrophil infiltration (Fig. 1E). M2 to M1 polarization was evidenced by the results that the number of inflammatory monocyte-derived macrophages (CCR2^+^) and M1 macrophages/dendritic cells (CD11C^+^) was significantly higher in the livers of *Lep^∆I14/∆I14^* rats after week 16, while that of M2 macrophages (CD163^+^) was less than the controls (Fig. 5D, fig. S5). The temporal examination of immune cell infiltration from week 4 to week 48 is included in fig. S5. As stated above, such phenotypes were not observed in *Leptin* deficiency mice raised with the same condition (Fig. 2, fig. S2). Taken together, hepatic inflammation in *Lep^∆I14/∆I14^* rats is evoked but restricted at certain level since week 16, which results from combinatory effects of gene expression, inflammatory signaling pathways and immune cell infiltration.

**Fig. 5.**
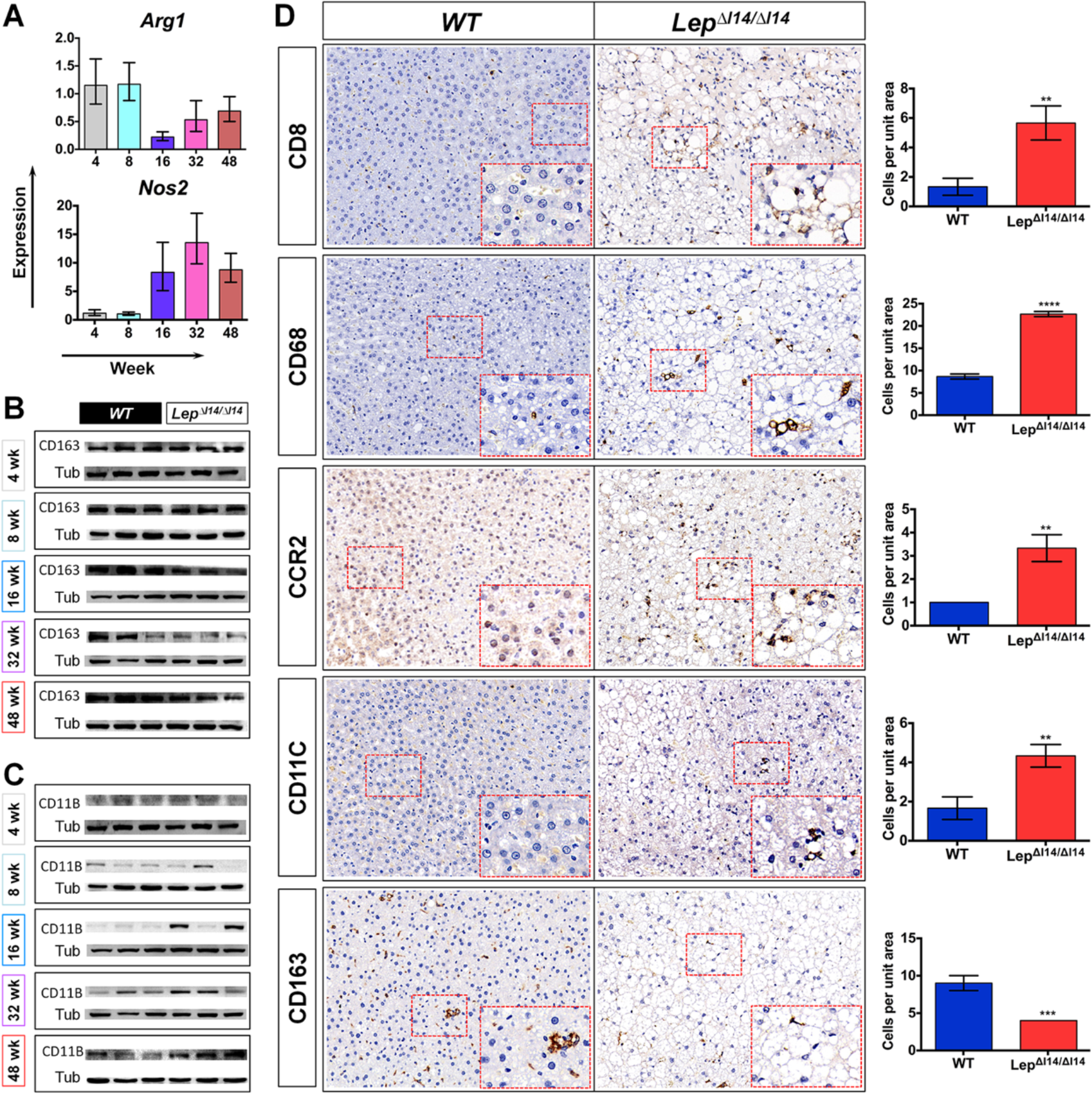
Immune cell infiltration during NASH progression in *Lep^∆I14/∆I14^* rats. (A) Dynamic changes of *Arg1* and *Nos2* expression verified by qRT-PCR (n=3). (B) and (C) Western bolt shows the expression of CD163 (B) and CD11B (C) (n=3). (D) IHC of liver sections shows infiltration of different immune cells at week 16 in *Lep^∆I14/∆I14^* rats compared to the WT controls. Right panel shows the statistics of immune cell infiltration (n=3). Data are presented as Mean±SD. **p<0.01, ***p<0.001, ****p<0.0001.

### *Lep^∆I14/∆I14^* rats share more homologous genes with NASH patients than previously established mouse models

As stated above, *Lep^∆I14/∆I14^* rats develop NASH with a unique progression pattern. However, is the molecular etiology of NASH in *Lep^∆I14/∆I14^* rats evolutionally conserved and similar to that in human beings? To answer this question, we compared the DEGs in *Lep^∆I14/∆I14^* rats across stages (Fig. 3B) to those DEGs in human beings with healthy obesity (HO), NAFLD, and NASH from the previous study.(26) There were 42 DEGs that were significantly and concordantly regulated in both *Lep^∆I14/∆I14^* rats at week 16 and human NASH (Fig. 6A). In contrast, there were maximal 18 DEGs in the mouse models (26). This indicates that *Lep^∆I14/∆I14^* rats were better than mouse models to match human NASH at the molecular level of gene expression. Of note, none of these DEGs were common across all the time points (fig. S6). The most overlapped DEGs include lipid metabolism related genes such as *Fabp4* and *Fasn*, and inflammation related genes such as *Inhbe* (fig. S6). We next examined the hepatic gene expression in *Lep^∆I14/∆I14^* rats and mouse models at the functional level. We identified differentially regulated pathways by GSEA analysis (27) in *Lep^∆I14/∆I14^* rats and mouse models, respectively. Comparison analysis found that 20 pathways were significantly and concordantly regulated in both *Lep^∆I14/∆I14^* rats at week 16 and human NASH (Fig. 6B). In contrast, there were 24 in WTD mice which share the most pathways with human NASH (26). This implies that both *Lep^∆I14/∆I14^* rats and WTD mice similarly match human NASH at the functional level. PharmaLegacy Laboratories (Shanghai) has initially screened 41 crab-eating monkeys for spontaneous NAFLD models (36 NAFLD and 5 healthy controls). Although these 36 NAFLD monkeys showed mild NAFL phenotypes with higher body weight, liver triglyceride, and histological simple steatosis, there was no significant difference in inflammation-related serum parameters (e.g., AST and ALT) or histological fibrosis compared to the controls (data not shown). So we wondered if transcriptome analysis can distinguish the monkeys’ mild NAFL phenotype from that of the severe NASH patients. We randomly chose 3 NAFLD and 3 control monkeys and analyzed their hepatic gene expression by RNA-seq (Their histological examination are shown in fig. S7, and serum parameters in table S5.). We found that the NAFLD monkeys shared as few as 5 DEGs (up-regulated *Fasn* and *Kcnj3*; down-regulated *Hal*, *P4ha1* and *Slc16a7*) with NASH patients, which was consistent with their mild NAFL phenotypes as described above. In addition, this suggests that a suitable NASH model is not determined by how closely the chosen species is related to human beings, but rather determined by whether it can faithfully model NASH progression in human beings to the greatest extent. Taken together, *Lep^∆I14/∆I14^* rats can serve as a useful NASH model as their hepatic transcriptome reflects NASH progression in humans more closely than other reported rodent models.

**Fig. 6.**
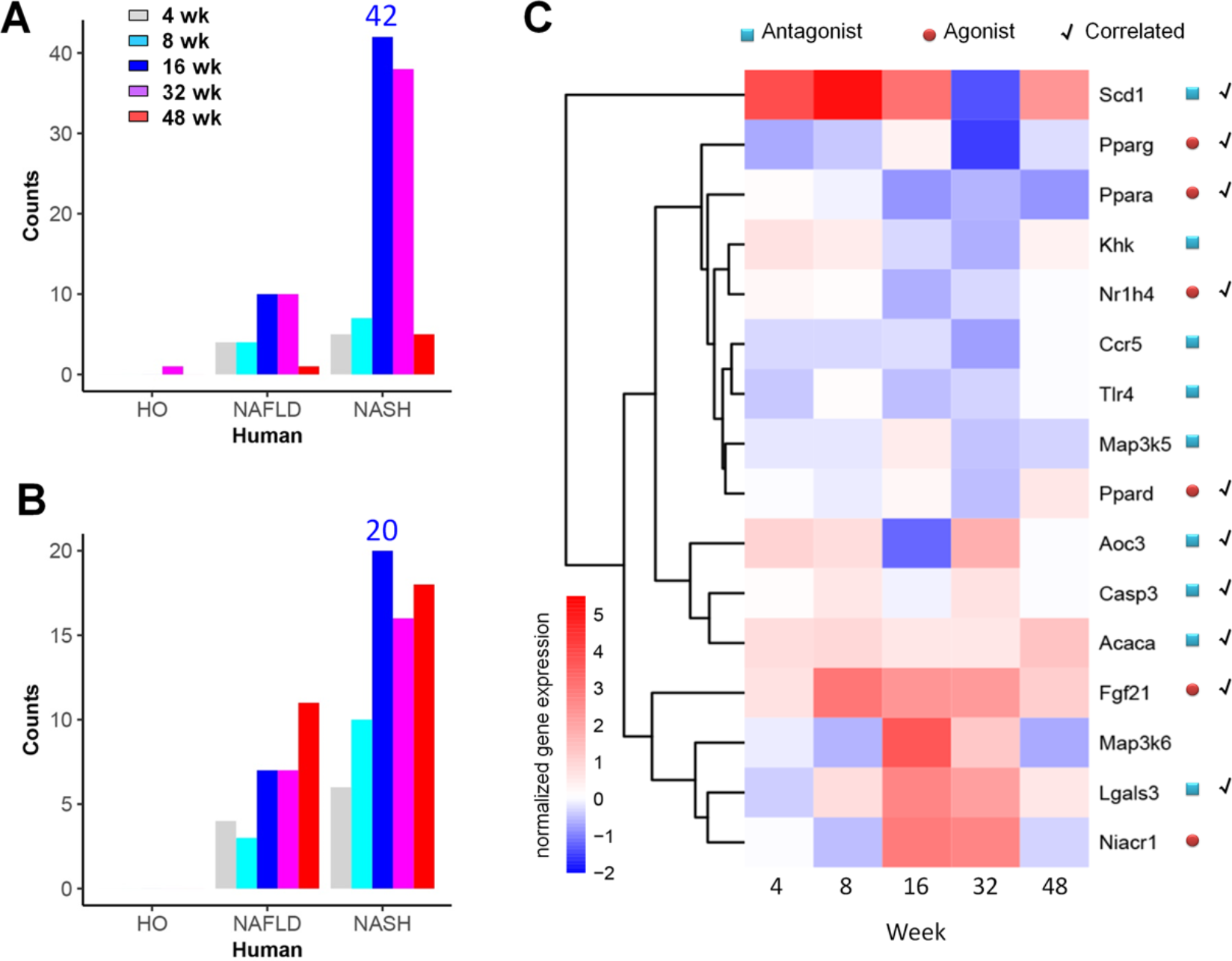
NASH progression in *Lep^∆I14/∆I14^* rats are evolutionarily conserved to human beings. (A) and (B) Counts of the significantly differentially regulated genes (A) and pathways (B) in *Lep^∆I14/∆I14^* rats that are also significantly and concordantly regulated in human heathy obese (HO), NAFLD and NASH, respectively. (C) Heatmap showing expression levels of the drug targets in clinical trials. Blue squires indicate antagonist targets; Red circles indicate agonist targets; Check marks indicate that gene expression is correlated with the targeted drug effect.

### *Lep^∆I14/∆I14^* rats have potential to evaluate drug targets at transcriptional level

Many drugs for potential treatment of NASH are in clinical trials, although none have been approved by FDA yet.(28, 29) So we asked if *Lep^∆I14/∆I14^* rats can be used to evaluate the drug targets, at least those in clinical trials. In the hepatic targets (28, 29) excluding non-detected *Ccr2* and *Sirt1*, agonist targeting genes such as *Fxr (Nr1h4)* and *Pparα /δ /γ* were down-regulated at most of the time points while antagonist targeting genes such as *Scd1*, *Acc (Acaca)*, *Aoc3*, *Casp3* and *Galectin3(Lgals3)* were up-regulated (Fig. 6C). *Fgf21* that encodes an endocrine sensor for nutrition context is upregulated as previously reported (30). This suggests that the pharmacologic agents targeting these genes may be pre-clinically tested in *Lep^∆I14/∆I14^* rats. Some targets could not be tested in *Lep^∆I14/∆I14^* rats. These targets included *Ccr5*, *Khk*, and *Hcar2 (Niacr1)* which did not show expected correlation. *Tlr4* mRNA level did not correlate well with its protein level (fig. S4C). *Ask2 (Map3k6)* instead of *Ask1 (Map3k5)* seemed to be up-regulated in *Lep^∆I14/∆I14^* rats (Fig. 6C). Nevertheless, *Lep^∆I14/∆I14^* rats are suitable for evaluation of many drug targets at transcriptional level and thus hold promise for screening potential biomarkers and treatment of NASH.

## Discussion

We discovered *Lep^∆I14/∆I14^* rats as a new animal model to study human NASH. For the past decades, many animal models have been created to study NASH.(9–11, 15) Dietary induction models are widely used, but human beings rarely consume such experimental diet to develop NAFLD. Sometimes dietary induction models do not recapitulate the phenotypes in human diseases. For example, MCD mouse model shows low body weight with low serum INSULIN and LEPTIN levels.(31) Genetic models such as *ob/ob* and *db/db* mice develop simple steatosis with regular diet, however a secondary hit, e.g., MCD (13), is required to develop NASH phenotypes. Nowadays, the drug development usually targets specific molecules. The molecular etiology, especially the spatial and temporal gene expression, should be fully characterized in the animal models before the pre-clinical examination of potential drugs. Unfortunately, the temporal progression of the disease is still not well investigated in most of the animal models. *Lep^∆I14/∆I14^* rats develop spontaneous NASH phenotypes, which makes the experiment less time and labor consuming. Given that rats are widely used in pharmacology and toxicology (32), revealing the natural history of NASH in *Lep^∆I14/∆I14^* rats will greatly facilitate further usage of this model to screen new drugs and treatment for NASH.

There are many hypotheses on the natural history of NASH, including ‘‘two hit’’ model (33) and “the multiple parallel hits hypothesis” (34). Using *Lep^∆I14/∆I14^* rats, we are able to examine the natural history of NASH at transcriptional level (Fig. 7). Although we cannot exclude other pathways which may also contribute, more or less, to NASH progression, we focused on distinct gene expression patterns in two major pathways such as lipid metabolism and inflammation. In *Lep^∆I14/∆I14^* rats, TAG accumulation in liver is correlated with increase in expression of genes encoding rate-limiting enzymes in lipid metabolism instead of the entire fatty acid biosynthesis (Fig. 7). Interestingly, the expression of hepatic inflammation related genes does not increase rectilinearly but is elevated and maintained at a high level after week 16, which is associated with the activation of JNK and NFκB pathways and immune cell infiltration (Fig. 7). The present work focuses on hepatic gene expression, but we perceive that many other tissues and organs must be involved in hepatic phenotypes of NASH, which should be further addressed using appropriate models.

**Fig. 7.**
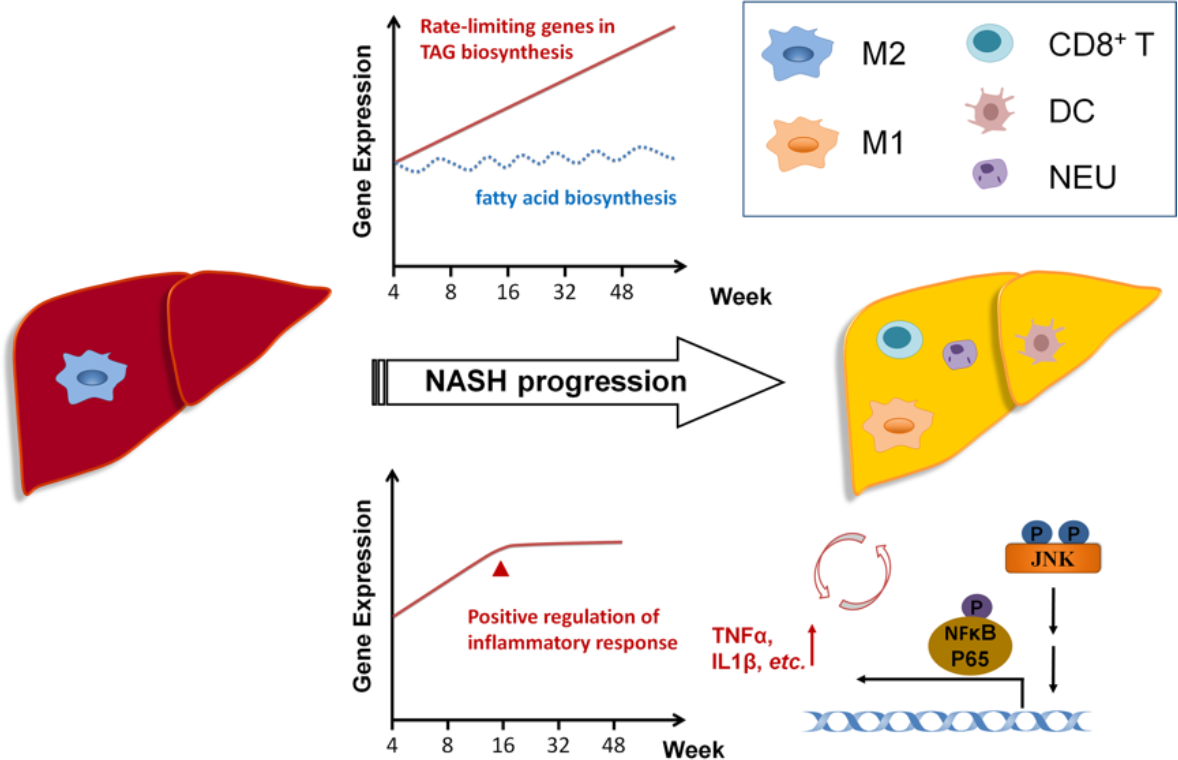
NASH progression in *Lep^∆I14/∆I14^* rats involves combinatory effects of gene expression, inflammatory signaling pathways and immune cell infiltration. During NASH progression in *Lep^∆I14/∆I14^* rats, TAG biosynthesis is associated with increased expression of the genes encoding rate-limiting enzymes, but not that of all the genes in fatty acid biosynthetic process. Meanwhile, the hepatic inflammation related genes such as *Tnfα* and *IL1β* are upregulated at week 16 (arrow) and then maintained at a high level till week 48. This transcriptional alteration is associated with the activation of JNK and NFκB pathways and immune cell infiltration, e.g. neutrophils, CD8^+^ T cells, and macrophages. TAG, triglyceride; M1, classically activated macrophages; M2, alternatively activated macrophages; CD8^+^ T, CD8^+^ T cells; DC, dendritic cells; Neu, neutrophils.

As a pro-inflammatory factor, LEPTIN plays essential roles in regulation of hepatic inflammation. Neither *ob/ob* mice nor *Lep^∆I14/∆I14^* rats develop fibrosis, since LEPTIN is required for the development of fibrosis in NASH (13), which is likely associated with activation of Stellate cells (35) and up-regulation of norepinephrine (36). However, hepatic inflammation in *Lep^∆I14/∆I14^* rats is kept higher than their mouse counterparts probably due to species difference. Previous research observed that LEPTIN resistance is present in NAFLD patients, however serum LEPTIN level does not correlate with NASH progression.(37) Indeed, our data suggest that the impairment of LEPTIN signaling maintains a restricted level of hepatic inflammation, which may also explain why only about 20% NAFLD patients develop NASH.(5) (28) LEPTIN signaling pathway may serve as an evolutionary protection mechanism against tissue damage from inflammation when excess nutrient intake is inevitable and irreversible. Therefore, we should carefully consider overdiagnosis/overtreatment of NAFLD as pointed out by Rowe (5) recently.

## Acknowledgments

The authors thank PharmaLegacy Laboratories (Shanghai) Co., Ltd for the generosity of providing the samples of *Macaca fascicularis* with spontaneous NAFLD. The authors thank Dr. Felice Elefant at Drexel University for revising the text. The work was supported by National Program on Key Basic Research Project (973 Program 2015CB964702, 2015CB964601); National Natural Science Foundation of China (81570521, Key Program 81430026); Experimental Animal Research Fund, and Science and Technology Commission of Shanghai Municipality (15140903900).The authors thank PharmaLegacy Laboratories (Shanghai) Co., Ltd for the generosity of providing the samples of *Macaca fascicularis* with spontaneous NAFLD; The authors thank Dr. Felice Elefant at Drexel University for revising the text.

## List of Abbreviations

ALT: alanine aminotransferase
AST: aspartate aminotransferase
DEGs: differentially expressed genes
GSEA: Gene Set Enrichment Analysis
GO: gene ontology
HE: hematoxylin and eosin
HCC: hepatocellular carcinoma
HDL-C: High Density Lipoprotein cholesterol
HO: healthy obese individuals
LDL-C: Low Density Lipoprotein cholesterol
LR: Leptin Receptor
MRE: magnetic resonance elastography
MRI-PDFF: magnetic resonance imaging-proton density fat fraction
NAFL: nonalcoholic fatty liver
NAFLD: nonalcoholic fatty liver disease
NASH: nonalcoholic steatohepatitis
TAG: triglyceride
WT: wild-type

## Supplementary data

**Fig. S1.**
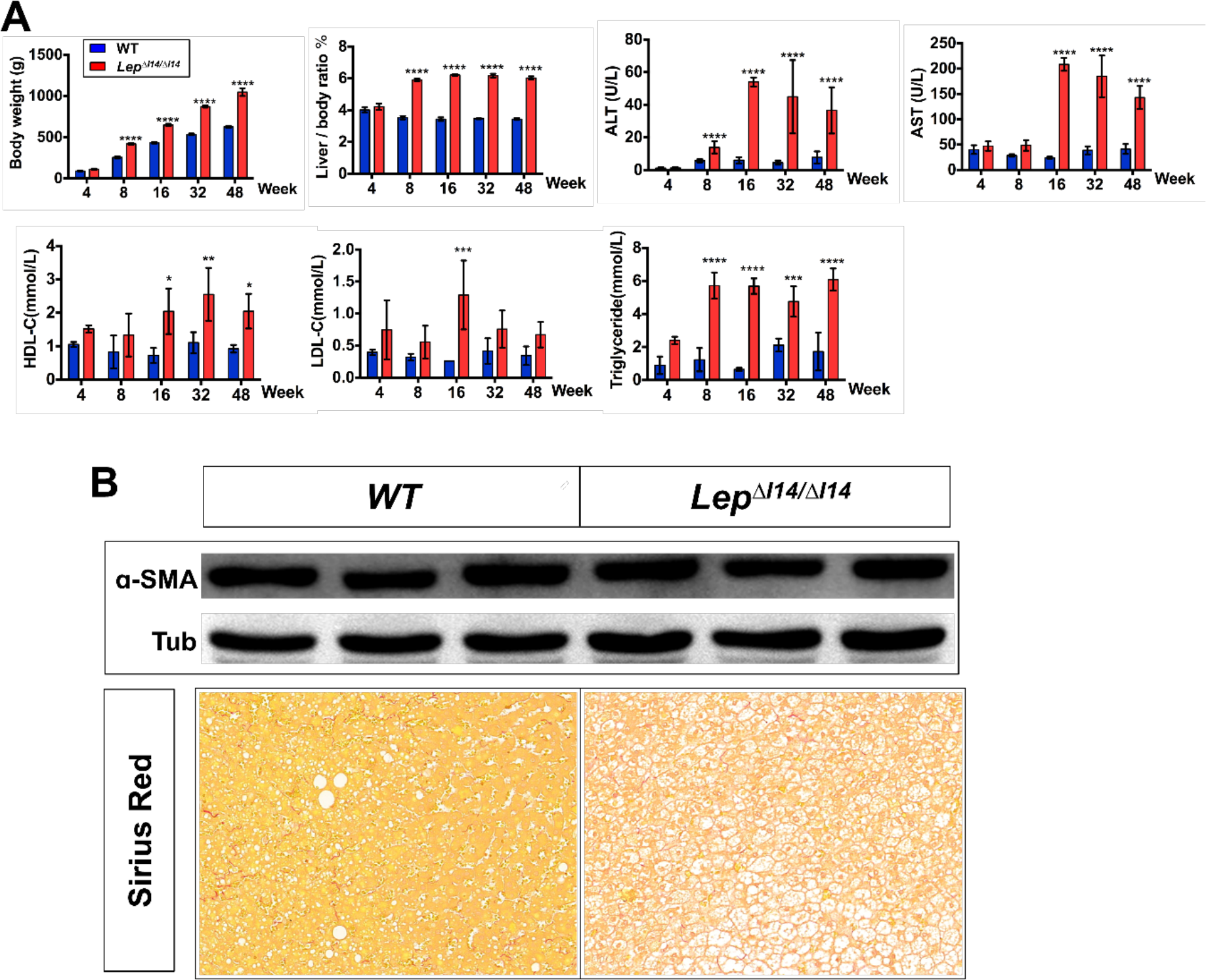
NASH phenotypes of *Lep^∆I14/∆I14^* rats. (A) Body weight, liver/body ratio, ALT and AST in *Lep^∆I14/∆I14^* rats from postnatal week 4 to 48, compared to the WT controls (n=6). Serum triglyceride, HDL-C and LDL-C levels in *Lep^∆I14/∆I14^* rats (n=3). Data are presented as Mean±SD. *p<0.05, **p<0.01, ***p<0.001, ****p<0.0001. (B) Representative data at week 16 showing that *Lep^∆I14/∆I14^* rats do not develop liver fibrosis. Western blot showed that a-SMA expression is not increased in *Lep^∆I14/∆I14^* rats. Compared to the WT controls, fibrosis is not seen in the liver sections by Sirius red staining.

**Fig. S2.**
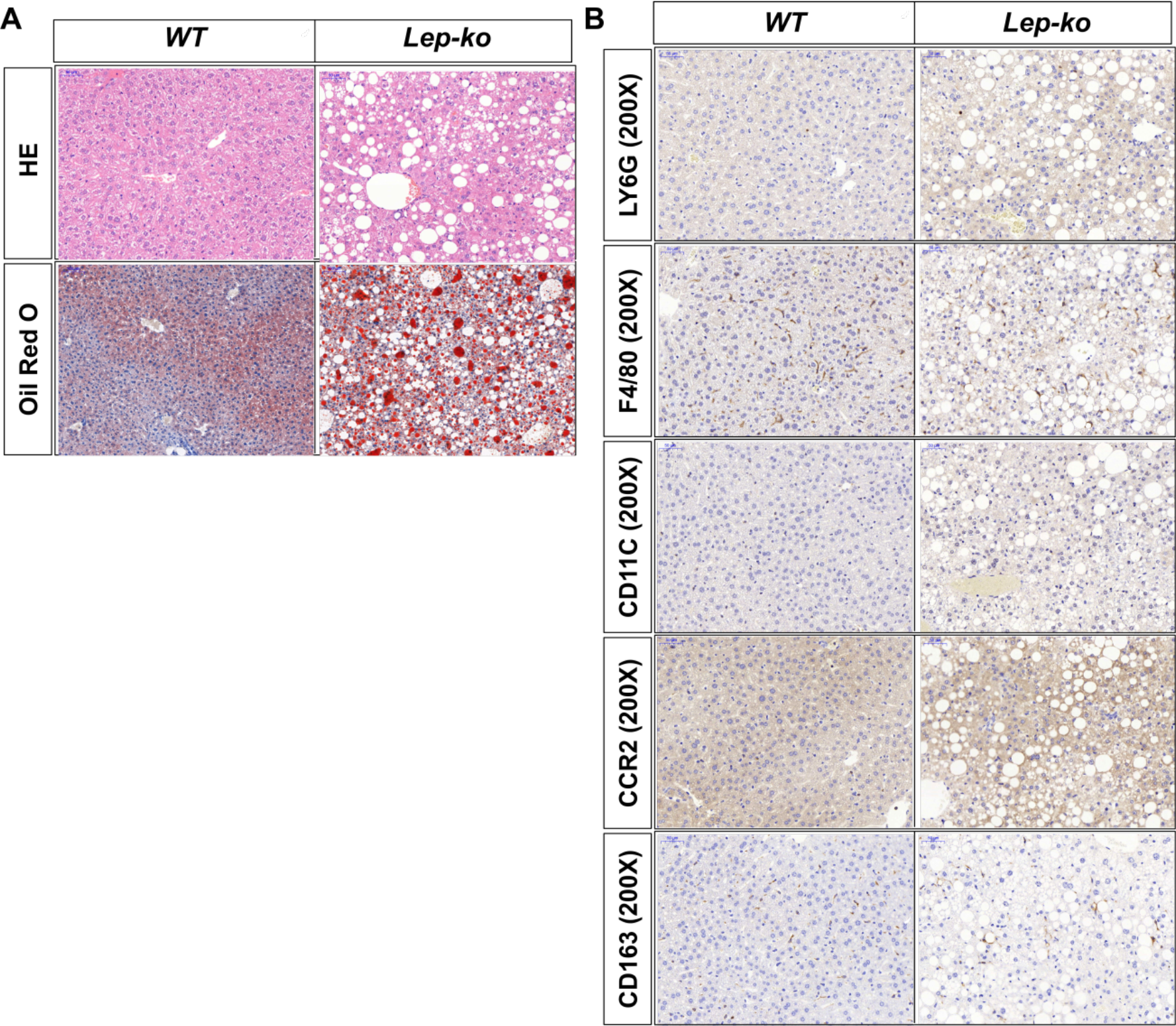
*ob/ob* mice do not develop NASH phenotypes at week 32. (A) HE and Oil red O staining showed liver steatosis at week 32. (B) Immune cell infiltration in liver showed no difference between *ob/ob* mice and the WT controls at week 32.

**Fig. S3.**
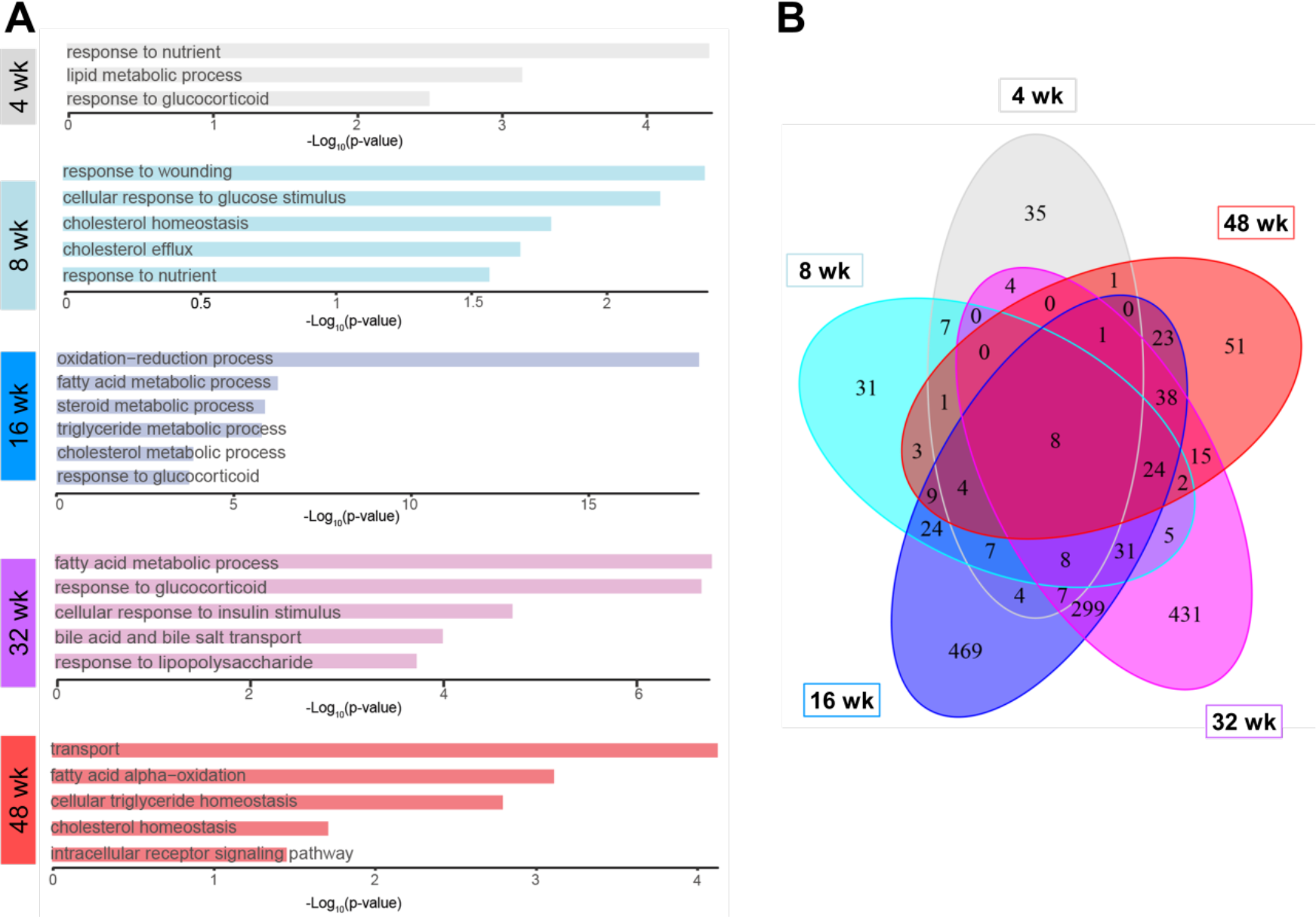
Characteristics of the down-regulated DEGs. (A) Down-regulated GO of DEGs at different time points. (B) Venn plot shows the overlap of down-regulated DEGs among different time points.

**Fig. S4.**
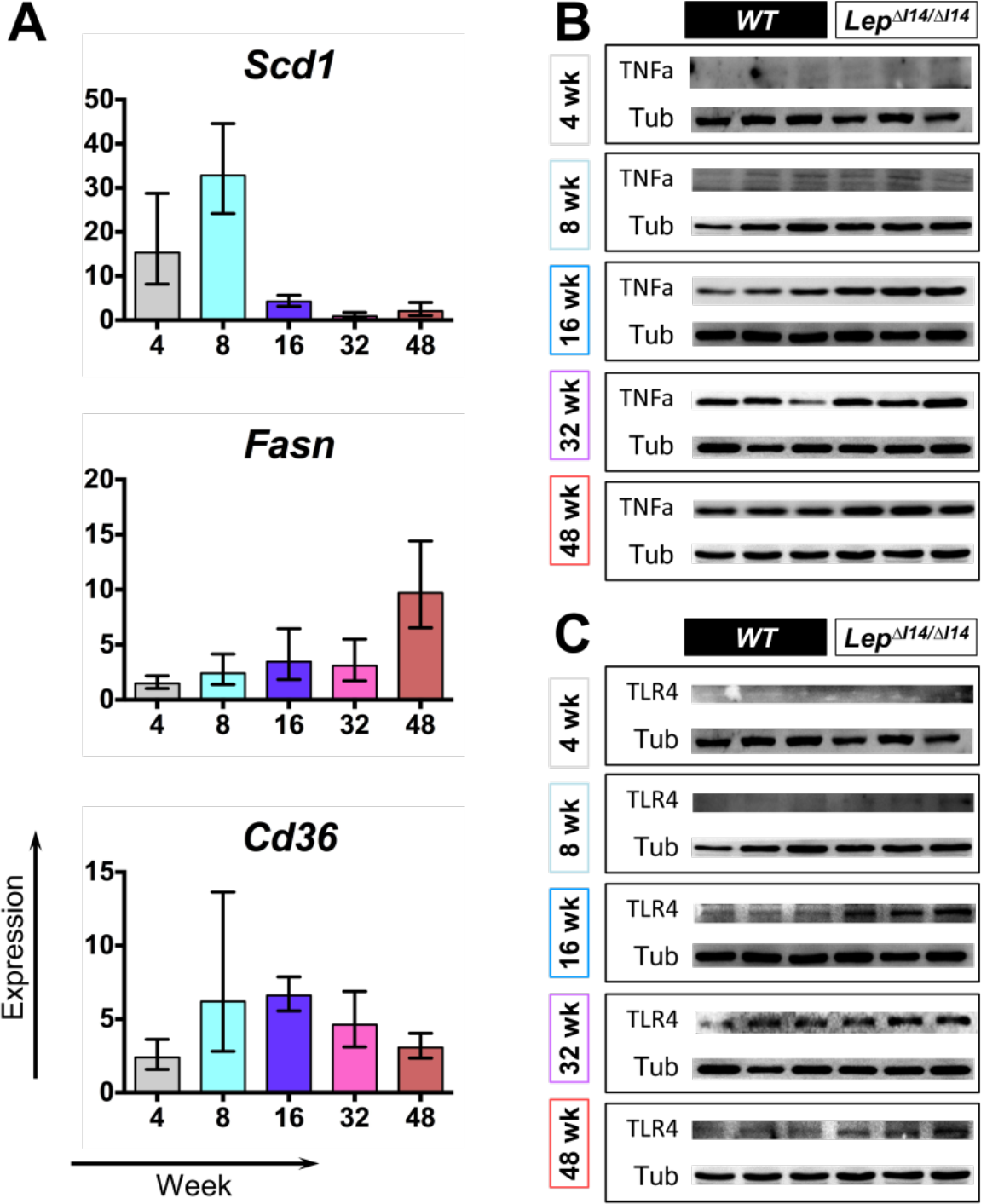
Gene expression verified by qPCR and Western blot. (A) Relative expression of *Cd36*, *Fasn* and *Scd1* using *Actb* as internal control. (B) and (C) Western blot shows the expression levels of phosphorylated Tnfα (B) and TLR4 (C) at different time points (n=3). β-TUBULIN is referred to as internal control.

**Fig. S5.**
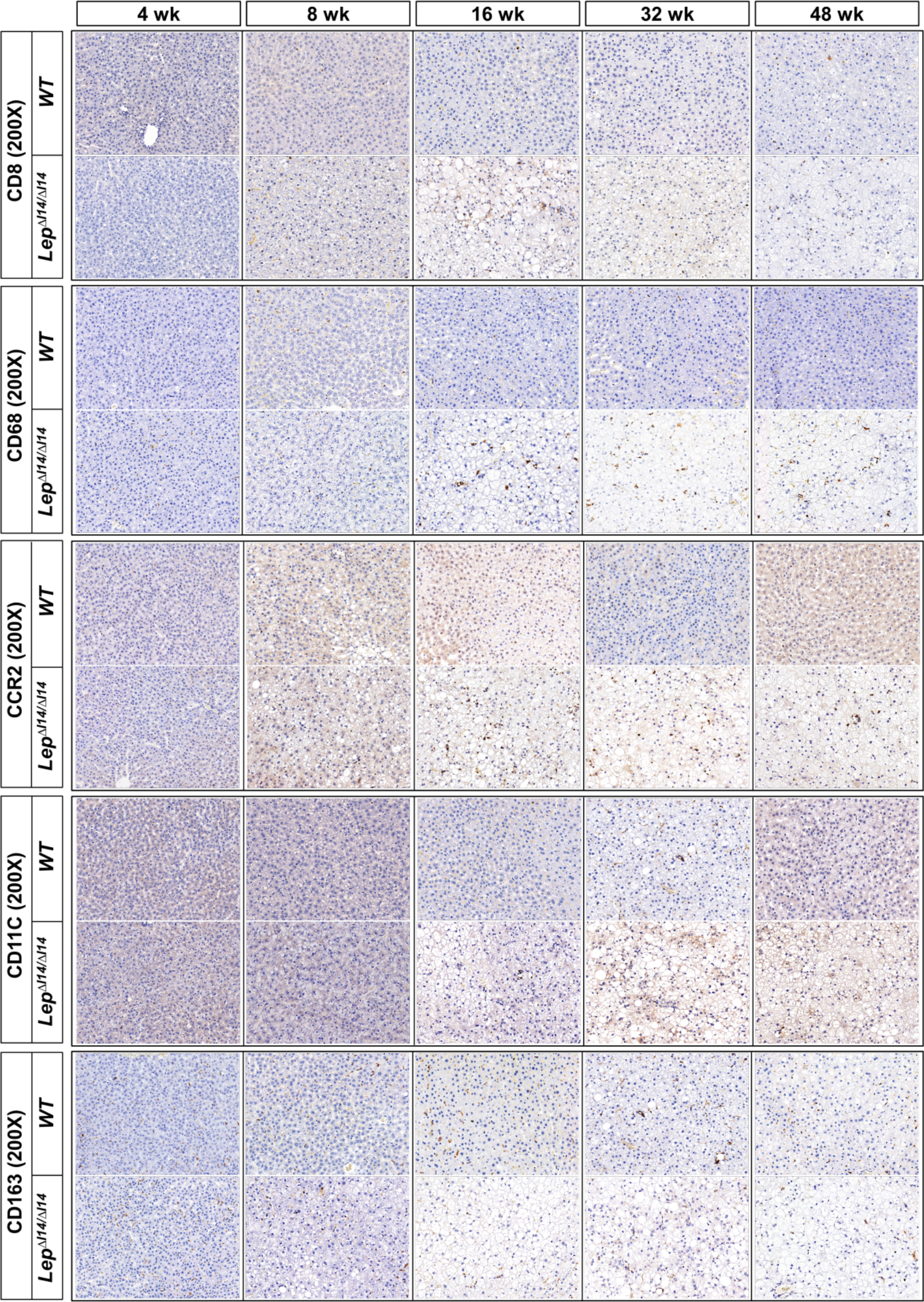
Representative IHC of liver sections shows immune cell infiltration at different time points during NASH progression.

**Fig. S6.**
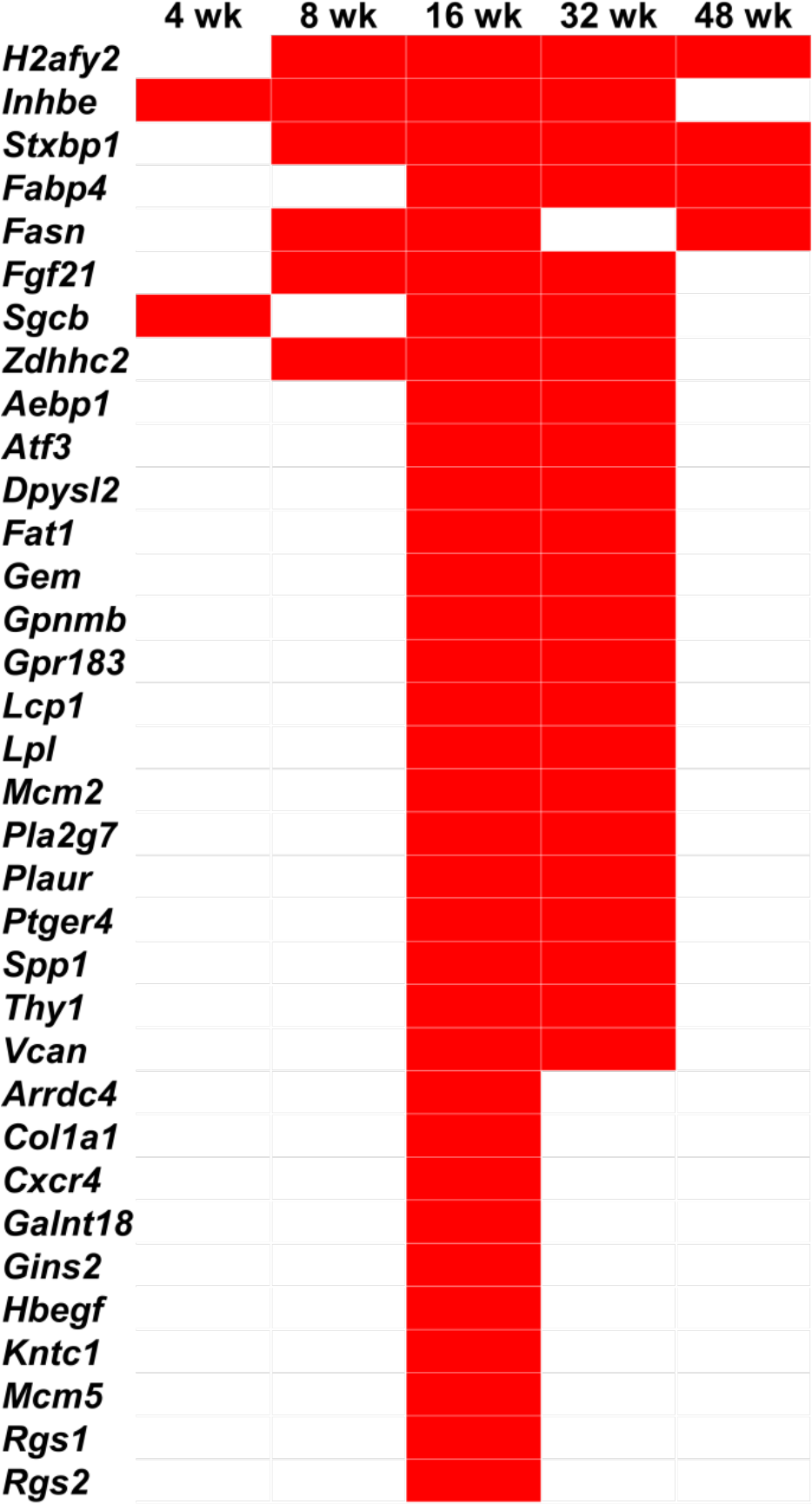
The DEGs that *Lep^∆I14/∆I14^* rats shared with NASH patients. Red bars indicate their overlap among different time points.

**Fig. S7.**
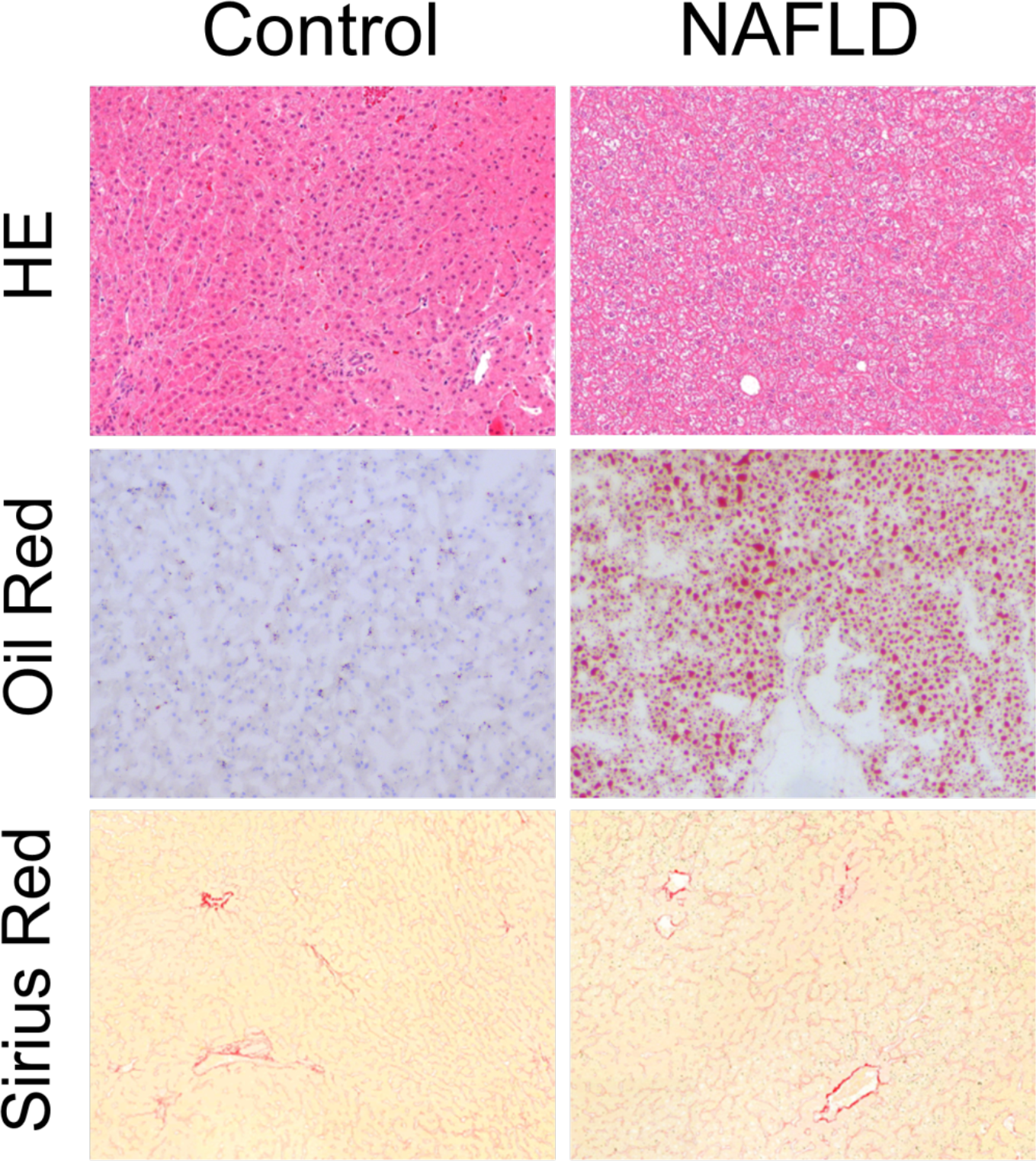
Compared to the controls, the obese crab-eating monkeys have hepatic steatosis, evidenced by liver biopsy histology (HE and Oil red O staining). However, they do not have liver fibrosis (Sirius red).

**Table S1.**
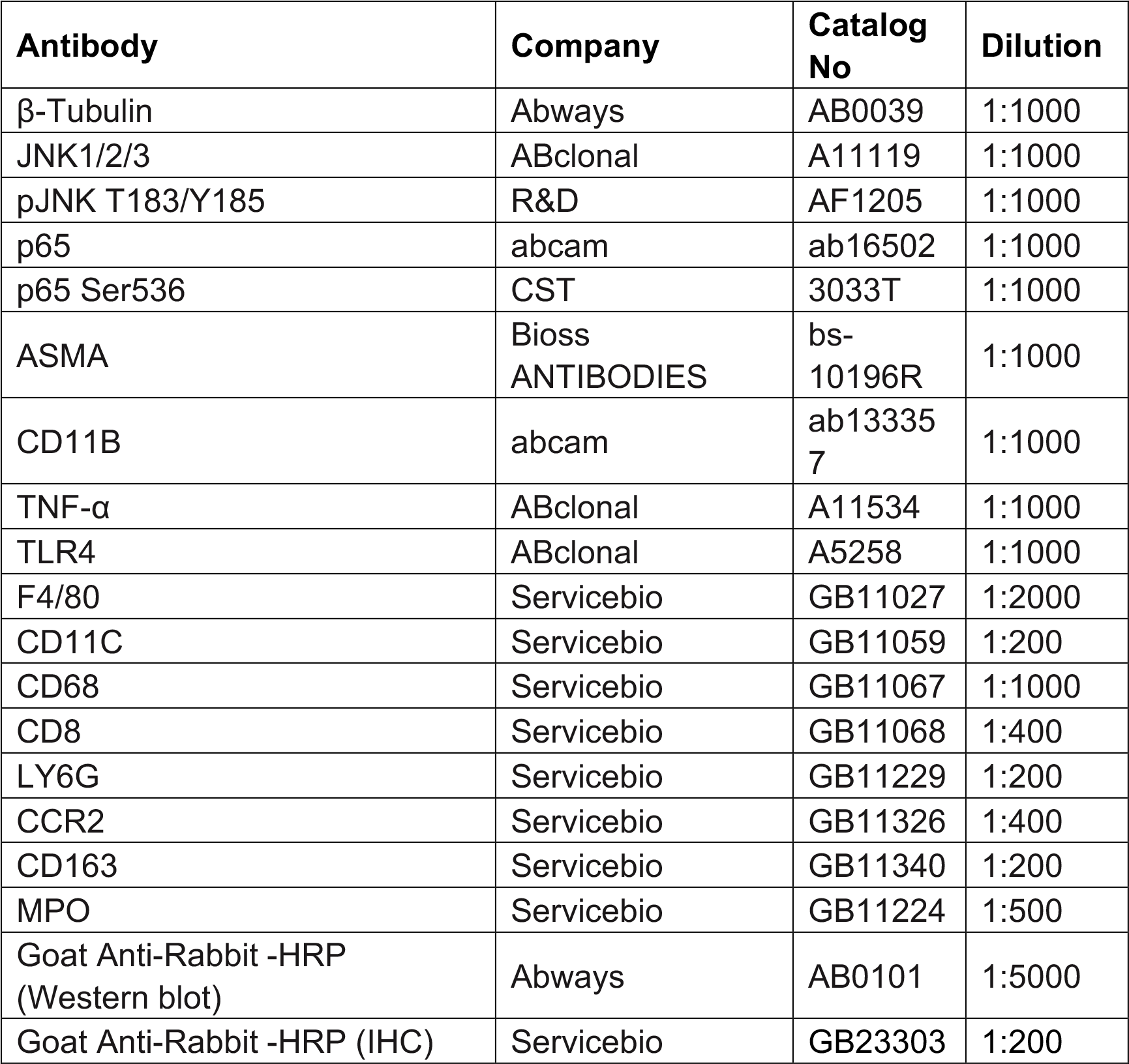
The antibody information used in IHC and Western blot.

**Table S2.**
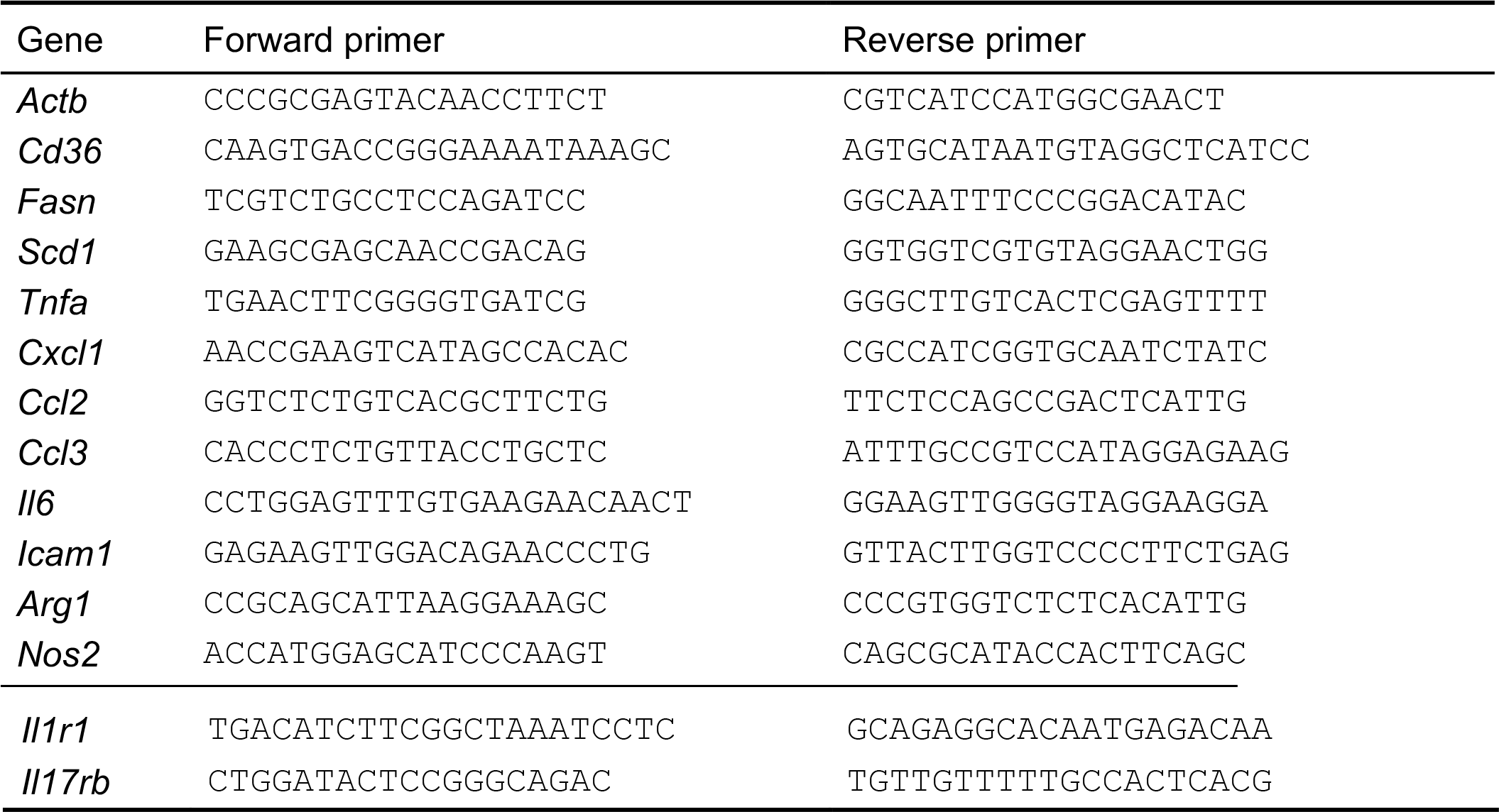
qPCR primer information.

**Table S3.** NASH scoring system used to evaluate the histology of rat livers.

| Criteria | Definition | Score |
| --- | --- | --- |
| Steatosis grade | <5% | 0 |
|  | 5%–33% | 1 |
|  | 33%–66% | 2 |
|  | >66% | 3 |
| Lobular inflammation | No foci | 0 |
|  | <2 foci per 200X field | 1 |
|  | 2–4 foci per 200X field | 2 |
|  | >4 foci per 200X field | 3 |
| Ballooning | None | 0 |
|  | Few balloon cells | 1 |
|  | Many cells/prominent ballooning | 2 |

**Table S4.** The DEGs shared among different time points.

|  | Up-regulated DEGs |  |  |  |  |  | Down-regulated DEGs |  |  |  |  |
| --- | --- | --- | --- | --- | --- | --- | --- | --- | --- | --- | --- |
|  | 4 wk | 8 wk | 16 wk | 32 wk | 48 wk |  | 4 wk | 8 wk | 16 wk | 32 wk | 48 wk |
| 4 wk | 195 | 56 | 56 | 35 | 23 |  | 88 | 36 | 39 | 29 | 16 |
| 8 wk | 56 | 249 | 140 | 70 | 63 |  | 36 | 165 | 115 | 79 | 52 |
| 16 wk | 56 | 140 | 1682 | 794 | 113 |  | 39 | 115 | 956 | 416 | 107 |
| 32 wk | 35 | 70 | 794 | 1366 | 85 |  | 29 | 79 | 416 | 874 | 89 |
| 48 wk | 23 | 63 | 113 | 85 | 198 |  | 16 | 52 | 107 | 89 | 181 |

**Table S5.** The histological examination and serum parameters of the spontaneous NAFLD monkey model.

| Item | WT | FL | Unit | P Value |
| --- | --- | --- | --- | --- |
| GLU | 1.797 ± 0.08838 | 2.067 ± 0.2858 | mmol/L | 0.4178 |
| TBL | 3.667 ± 0.2963 | 1.967 ± 0.1453 | μmol/L | 0.0067 |
| DBL | 0.5667 ± 0.1202 | 0.2667 ± 0.03333 | μmol/L | 0.0739 |
| IBIL | 3.100 ± 0.2000 | 1.700 ± 0.1155 | μmol/L | 0.0037 |
| TP | 70.93 ± 0.6333 | 62.13 ± 0.4842 | g/L | 0.0004 |
| ALB | 40.13 ± 1.035 | 34.83 ± 1.481 | g/L | 0.0427 |
| GLB | 30.80 ± 1.012 | 27.30 ± 1.200 | g/L | 0.0896 |
| A/G | 1.333 ± 0.08819 | 1.300 ± 0.1000 |  | 0.8149 |
| AST | 59.00 ± 2.887 | 51.33 ± 6.173 | U/L | 0.3235 |
| ALT | 57.00 ± 18.88 | 29.33 ± 6.227 | U/L | 0.2364 |
| AST/ALT | 1.240 ± 0.3355 | 1.957 ± 0.5051 |  | 0.3027 |
| ALP | 74.67 ± 1.764 | 70.00 ± 14.80 | U/L | 0.7698 |
| R-GT | 62.00 ± 11.02 | 62.00 ± 10.60 | U/L | > 0.9999 |
| AFU |  | N/A | U/L | N/A |
| TBA | 2.467 ± 1.220 | 3.900 ± 1.823 | μmol/L | 0.5491 |
| GPDA | 118.0 ± 22.11 | 86.00 ± 19.08 | U/L | 0.3348 |
| TG | 0.4167 ± 0.1368 | 0.2333 ± 0.04910 | mmol/L | 0.2757 |
| T-CH | 1.817 ± 0.1670 | 2.250 ± 0.04933 | mmol/L | 0.0676 |
| HDL-C | 0.8500 ± 0.06028 | 1.090 ± 0.05508 | mmol/L | 0.0424 |
| LDL-C | 0.9067 ± 0.1424 | 0.9867 ± 0.01856 | mmol/L | 0.6072 |
| INS | 91.90 ± 33.62 | 173.7 ± 94.74 | pmol/L | 0.4615 |
| APOE | 9.733 ± 0.7860 | 10.47 ± 0.2028 | mg/L | 0.4174 |

